# PD-L1:CD80 Heterodimer Triggers CD28 While Repressing Both PD-1 and CTLA4 Pathways

**DOI:** 10.1101/615138

**Authors:** Yunlong Zhao, Chia-Hao Lin, Calvin Lee, Xiaozheng Xu, Zhe Huang, Changchun Xiao, Jack Bui, Li-Fan Lu, Enfu Hui

## Abstract

Combined immunotherapy with anti-PD-1/PD-L1 and anti-CTLA4 has resulted in superior clinical responses compared to single agent therapy. The underlying mechanisms for this synergy have yet to be elucidated and investigations have largely focused on cellular interactions. Herein, we report a molecular crosstalk in which the PD-1 ligand PD-L1 and the CTLA4 ligand CD80 heterodimerize in *cis*. This heterodimerization blocks both PD-L1:PD-1 and CD80:CTLA4 interactions, but preserves the ability of CD80 to activate the T cell costimulatory receptor CD28. Remarkably, PD-L1 expression on antigen presenting cells (APCs) protects CD80 from CTLA4 mediated *trans*-endocytosis, and the therapeutic PD-L1 blockade antibody atezolizumab paradoxically downregulates CD80 on APCs, presumably reducing its co-stimulatory ability. Importantly, this effect can be negated by co-blockade of CTLA4 with ipilimumab. Our study reveals an unexpected immune stimulatory role of *cis*-acting PD-L1 and a mechanism of anti-PD-L1/anti-CTLA4 crosstalk, providing a therapeutic rationale for combination blockade of PD-L1 and CTLA4.

## INTRODUCTION

Programmed cell death-1 (PD-1) and cytotoxic T lymphocyte-associated protein-4 (CTLA4) are two central co-inhibitory receptors that restrict T cell activity (Nishimura et al., 1999; Tivol et al., 1995; Waterhouse et al., 1995). Antibodies that block CTLA4 or PD-1/PD-L1 pathways have demonstrated impressive clinical activities against an array of human cancers in a subset of patients (Hodi et al., 2010; Powles et al., 2014; Ribas and Wolchok, 2018; Rizvi et al., 2015; Topalian et al., 2012). Notably, the combination of anti-CTLA4 and anti-PD-1 has proven more effective than either agent alone and recently approved by the FDA for treatment of human melanoma and renal cancer patients (Callahan et al., 2014; Larkin et al., 2015; Wei et al., 2018). Nevertheless, durable response to this combination therapy is restricted to a minority of patients and cancer indications (Wei et al., 2018). Therefore, a better mechanistic understanding of anti-PD-1/PD-L1, anti-CTLA4, and their crosstalk, is needed for rational design of combination therapies.

Binding of T-cell-intrinsic PD-1 with its ligand PD-L1 (Dong et al., 1999; Freeman et al., 2000) on APCs triggers tyrosine phosphorylation of PD-1 and recruitment of SHP2, a phosphatase that dephosphorylates T cell receptor (TCR) and CD28 costimulatory signaling components (Hui et al., 2017; Parry et al., 2005; Sheppard et al., 2004; Yokosuka et al., 2012). CTLA4 outcompetes CD28 for their shared ligands, CD80 and CD86, due to its substantially higher affinities to both ligands (Krummel and Allison, 1995; Linsley et al., 1991; van der Merwe et al., 1997). Additionally, CTLA4 depletes CD80 and CD86 molecules from APCs via *trans*-endocytosis to indirectly inhibit CD28 signaling (Qureshi et al., 2011; Wing et al., 2008). Intriguingly, PD-L1 and CD80 were reported to bind each other, suggesting another layer of crosstalk among PD-1, CTLA4 and CD28 pathways. However, the biochemical nature and cell biology consequence of PD-L1:CD80 interaction remain elusive. The original reports suggest that PD-L1 and CD80 bind in *trans* from different cells (Butte et al., 2007; Butte et al., 2008), yet recent studies suggest that they interact in *cis* on the same cells (Chaudhri et al., 2018). Dissecting the function of PD-L1:CD80 interaction is challenging due to a complex network consisting of two ligands (PD-L1 and CD80) and three receptors (PD-1, CD28 and CTLA4). Here we employed in *vitro* reconstitution and engineered cell lines to decouple and dissect the five-protein signaling network. We elucidated the functional consequences of PD-L1:CD80 interaction and uncovered an unexpected crosstalk among CD28, PD-1 and CTLA4 pathways at the extracellular level.

## RESULTS

### PD-L1 and CD80 interact only in *cis*

We first asked whether PD-L1 binds to CD80 in *trans* from different membranes. To this end, we utilized a membrane adhesion assay (Zhao et al., 2018), in which *trans*-interaction between two proteins is measured as the association of two model membranes: large unilamellar vesicles (LUVs) and a supported lipid bilayer (SLB). Specifically, we reconstituted His-tagged ectodomains of PD-L1 (PD-L1-His) and its binding partner, CD80 or PD-1, to Bodipy-PE containing LUVs and an SLB, respectively, via the chelating lipid DGS-NTA-Ni. Incubation of PD-L1 LUVs with the PD-1 SLB led to a number of SLB-associated LUVs, as visualized by total internal reflection fluorescence (TIRF) microscopy. By contrast, CD80-SLB captured 99% fewer PD-1 LUVs, similar to SLB with SLB with CD86, a negative control with no reported PD-L1 binding activity (Figure 1A). Hence, PD-L1 does not bind CD80 in *trans*, consistent with a recent study(Chaudhri et al., 2018).

**Figure 1.**
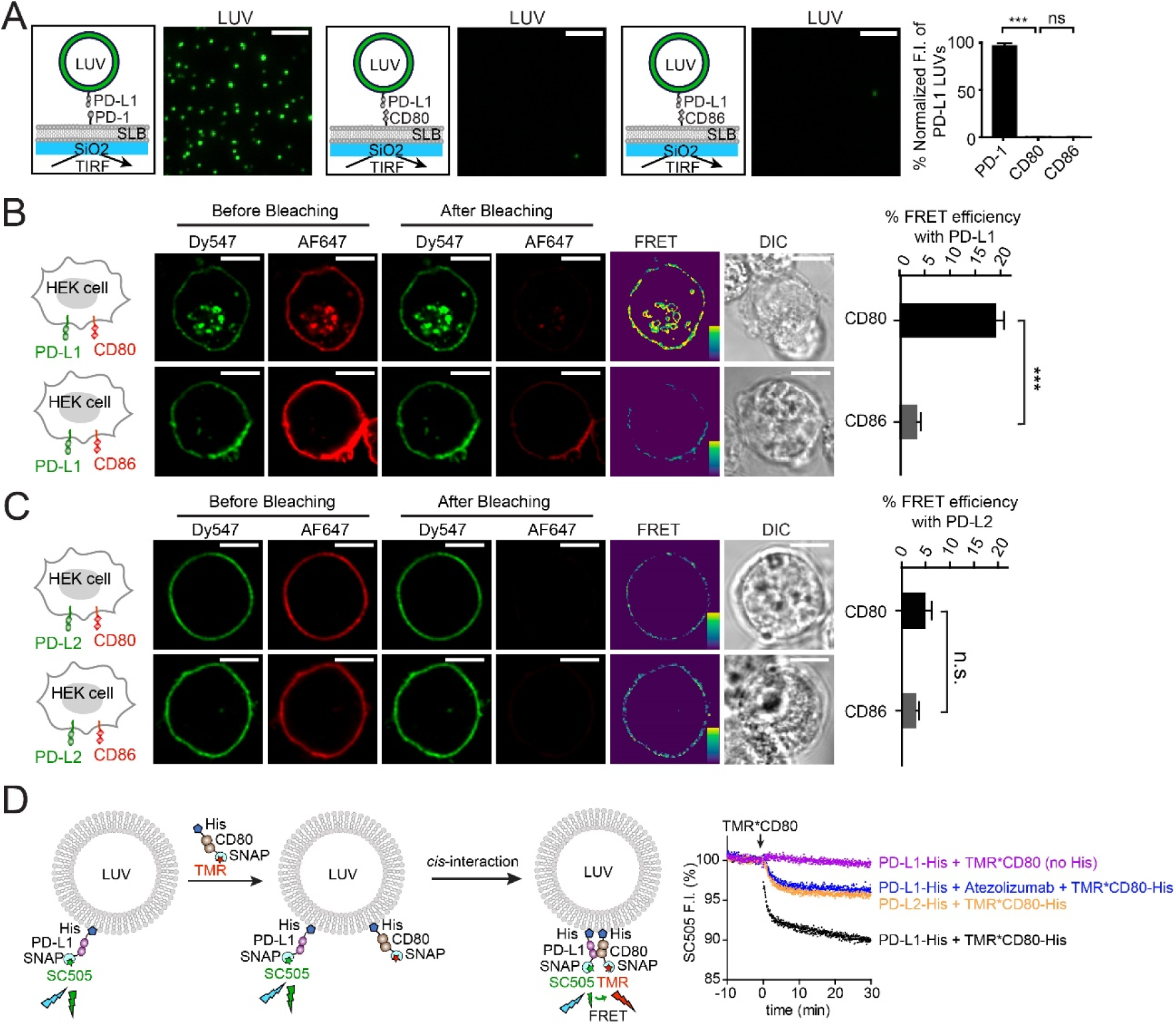
PD-L1 binds CD80 in *cis* and atezolizumab disrupts this interaction. **(A)** Representative TIRF images of PD-L1 LUVs captured by PD-1 SLB, CD80 SLB or CD86 SLB, with each LUV registered as a green dot. Bar graph summarizes the fluorescence intensity (F.I.) of the LUV channel under indicated condition, normalized to the intensity of the condition with PD-1 SLBs. Data are means ± SEM, n = 3. Scale bars: 5 μm. **(B)** An acceptor-photobleaching FRET assay showing PD-L1:CD80 *cis*-interaction on cell membranes. Cartoon on the left depicts a HEK293T cell co-expressing PD-L1 (labeled with Dy547, donor) and either CD80 or CD86 (labeled with AF647, acceptor). Pre- and post-bleaching confocal images are tabulated for a representative cell. Further right is the calculated FRET efficiency image (pseudo-color, with yellow to purple spectrum indicating strong to weak FRET, see **Methods**), and the differential interference contrast (DIC) image. Bar graph summarizes the FRET efficiencies as mean ± SEM, n > 25 cells from three independent experiments. Scale bars: 10 μm. **(C)** Same as **(A)** except replacing PD-L1 with PD-L2. **(D)** An LUV-reconstituted FRET assay for probing PD-L1:CD80 *cis*-interaction and Atetolizumab effect. As depicted in the cartoon on the left, SC505 (donor) labeled SNAP-PD-L1-His was pre-bound to LUVs via DGS-NTA-Ni lipid. Subsequently added TMR (acceptor) labeled SNAP-CD80-His bound to the LUVs, and interacts with PD-L1 in *cis*, causing FRET and SC505 quenching (black trace). Shown on the right are representative time courses of SC505 fluorescence under the indicated conditions. Data were normalized as described in **Methods**.

We next determined whether PD-L1 and CD80 bind in *cis* using a Förster resonance energy transfer (FRET) in a HEK293T cell reconstitution system (Zhao et al., 2018). To this end, we co-transfected HEK293T cells with CLIP-tagged PD-L1 and SNAP-tagged CD80, and labeled them with Dy547 (energy donor) and Alexa Fluor 647 (AF647, energy acceptor) respectively. Photo-bleaching of AF647-CD80 substantially increased the fluorescence of Dy547-PD-L1 (Figure 1B, **upper**), indicative of FRET. Replacement of CD80 with CD86 (Figure 1B, **lower**), or PD-L1 with PD-L2 dramatically decreased the FRET signal (Figure 1C). These data suggest that PD-L1 associates with CD80 in *cis* on cell membranes. We next examined this *cis*-interaction quantitatively in an LUV reconstitution assay. Specifically, we attached DGS-NTA-Ni containing LUVs with purified PD-L1-His labeled with energy donor [SNAP-Cell-505 (SC505)]. Subsequent addition of energy acceptor [tetramethyl rhodamine (TMR)] labeled CD80-His, but not CD80 lacking the membrane-targeting His-tag, quenched PD-L1 fluorescence (Figure 1D, black versus purple). This result indicates that PD-L1:CD80 *cis*-interaction on membranes has a much higher affinity than their interaction in solution (Cheng et al., 2013). CD80-His also induced a reproducible, but much weaker quenching of LUV-bound PD-L2 (Figure 1D, yellow), owing to a molecular crowding effect. Importantly, atezolizumab, a PD-L1 antibody that blocks both PD-L1:PD-1 interaction and PD-L1:CD80 interaction (Herbst et al., 2014), decreased the PD-L1:CD80 FRET to a similar level as PD-L2:CD80 FRET (Figure 1D, blue). These results demonstrate that PD-L1 and CD80 bind directly in *cis* and that this *cis*-interaction can be blocked by atezolizumab.

### PD-L1:CD80 *cis*-interaction blocks PD-L1:PD-1 signaling

Having established that CD80 binds PD-L1 in *cis*, we next sought to determine if CD80 inhibits PD-L1:PD-1 interaction. PD-1-Fc stained PD-L1 transduced, CD80 knockout Raji (PD-L1+/CD80 KO) B cells in a concentration dependent manner (Figure 2A, red). Co-expression of CD80 (3.5-fold excess of PD-L1) dramatically decreased PD-1-Fc staining (Figure 2A, blue, see Figure S1 for quantitation of PD-L1 and CD80 levels), indicating that PD-L1:CD80 *cis*-interaction inhibits PD-L1:PD-1 interaction. To extend the analysis to membrane-bound PD-1, we then employed a T cell–lipid bilayer hybrid system in which a cytotoxic T cell interacts with a SLB functionalized with peptide-linked-major-histocompatibility-complex (pMHC) and PD-L1. As reported (Hui et al., 2017; Yokosuka et al., 2012; Zhao et al., 2018), interaction of PD-1 on T cells with PD-L1 on SLB in *trans* led to the formation of PD-1 microclusters. Strikingly, addition of CD80-His (three-fold excess to PD-L1) to the PD-L1/pMHC SLB completely abolished PD-1 microclusters (Figure 2B). By contrast, equivalent levels of CD86-His did not affect PD-1 clustering (Figure 2B). TCR microclusters remained intact under all conditions (Figure S2). Thus, CD80 selectively inhibits PD-L1:PD-1 interaction.

**Figure 2.**
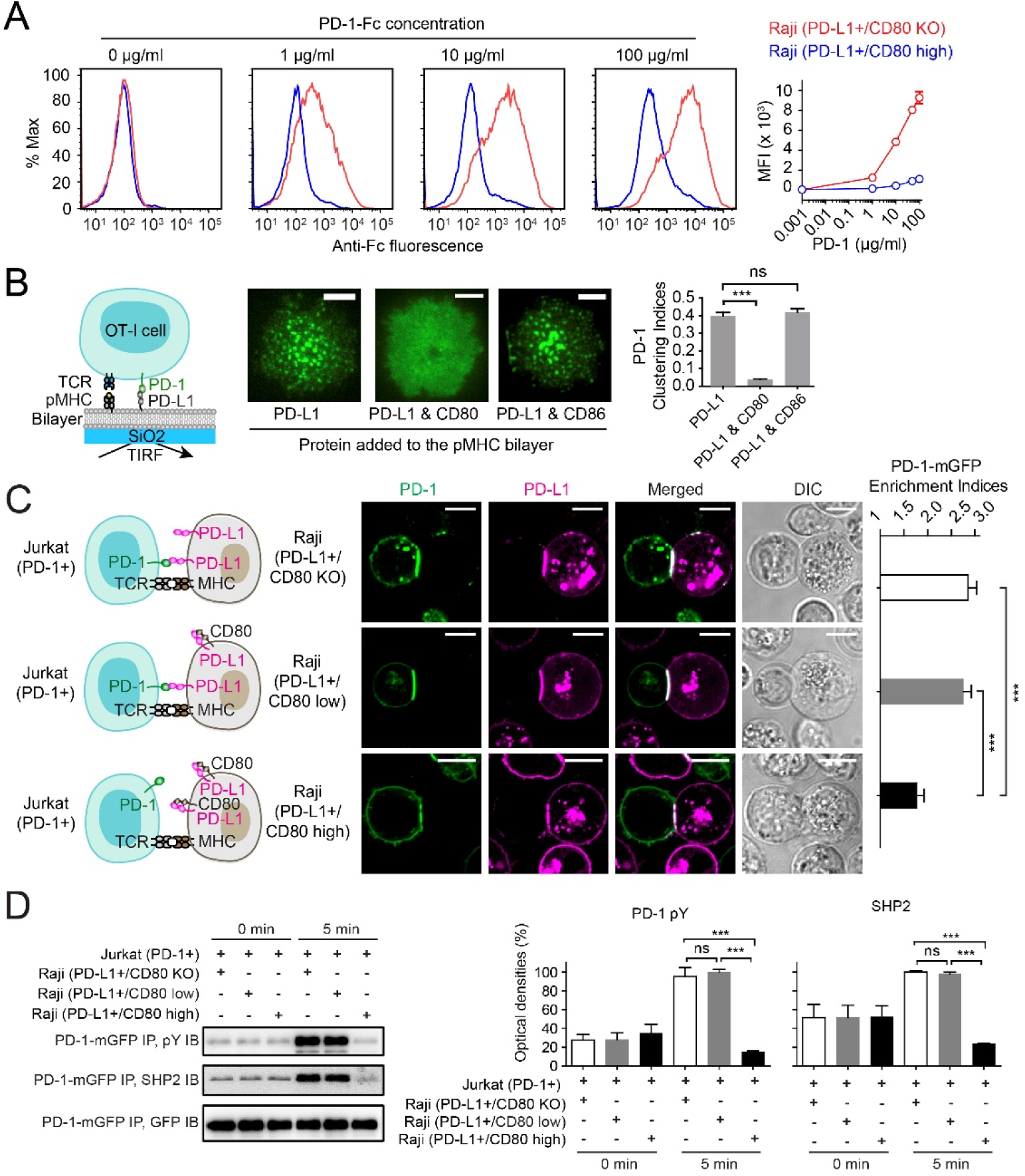
CD80 inhibits PD-1 signaling through neutralizing PD-L1 in *cis*. **(A)** Representative flow cytometry histograms of PD-1-Fc staining to the indicated types of Raji B cells at the indicated concentrations. Bound PD-1-Fc was labeled by anti-human IgG AF647 (see **Methods**). Mean fluorescence intensity (MFI) of AF647 was plotted against PD-1-Fc concentration (means ± SEM, n = 3). **(B)** A T cell–SLB assay for examining how CD80 affects the ability of PD-L1 to trigger PD-1 signaling clusters in cytotoxic T cells. As depicted in the cartoon, PD-1–mCherry transduced OT-1 cells interacts with SLB functionalized with pMHC and PD-L1. Shown are representative TIRF images of PD-1–mCherry (rendered in green) distribution 30 seconds after the OT-1 cell made contact with the SLB functionalized with peptide linked MHC class I (SIINFEKL-H2Kb, pMHC) plus PD-L1, PD-L1 & three-fold excess CD80, or PD-L1 & three-fold excess CD86. Bar graph on the right shows the clustering index of PD-1 in each condition. Data are the means ± SEM of at least 18 cells from three independent replicates. Scale bars: 5 μm. See Figure S1 for the corresponding images of TCR labeled by H57-597*AF647. **(C)** A T cell–APC conjugation assay showing CD80 on Raji B cells inhibits the synaptic enrichment of PD-1 from Jurkat T cells. Cartoons on the left depict PD-1– mGFP transduced Jurkat forming conjugate with three types of PD-L1–mCherry– transduced Raji B cells with increasing CD80 expression: CD80 KO Raji (PD-L1+/CD80 KO), Raji with endogenous level of CD80 (PD-L1+/CD80 low)), and CD80 overexpressing Raji (PD-L1+/CD80 high). Next to the cartoons are representative confocal images of the cell conjugate acquired two minutes after cell-cell contact. mGFP and mCherry signals are shown as green and magenta respectively. Scale bars: 10 µm. Bar graph on the right summarizes the synaptic enrichment indices (calculated as described in **Methods**) of the three conditions. Data are means ± SEM, n = 38 conjugates from three independent experiments. **(D)** Shown on the left are representative immunoblots (IB) showing the phosphorylation and SHP2 association of PD-1–mGFP, immunoprecipitated (IP) from the lysates of the indicated Jurkat-Raji co-culture. Cells were conjugated as described in **Methods**, and the times at which the co-cultures were lysed are indicated. Shown on the right are bar graphs summarizing the quantification of the blots (means ± SEM, n = 3). See also Figure S1, Figure S2, and Figure S3.

We next extended our investigation to a T cell-APC co-culture system consisting of PD-1–mGFP transduced Jurkat T cells and PD-L1-mCherry transduced Raji APCs (Tian et al., 2015). We created three Raji lines expressing similar levels of PD-L1-mCherry (~1700 molecules per μm^2^) but increasing levels of CD80: PD-L1+/CD80 KO), PD-L1+/CD80 low (~600 CD80 molecules per μm^2^), and PD-L1+/CD80 high (~6000 CD80 molecules per μm^2^) (Figure 2C & D, Figure S1). Notably, these cells express PD-L1 and CD80 at comparable levels with those on human monocytes derived dendritic cells (DCs) (Figure S3). Conjugation of superantigen SEE-loaded Raji (PD-L1+/CD80 KO) cells with Jurkat (PD-1+) cells enriched both PD-L1 and PD-1 to their interface. The CD80 low Raji cells, which express 2.9-fold lower CD80 than PD-L1 (Figure S1), had little effect on the PD-1 enrichment. The CD80 high Raji, which express ~3.5-fold higher CD80 than PD-L1, dramatically decreases PD-L1:PD-1 interface enrichment (Figure 2C), PD-1 phosphorylation and SHP2 recruitment (Figure 2D). Taken together, these results demonstrate that CD80 sequesters PD-L1 in *cis* to block PD-L1:PD-1 signaling, explaining a prior finding that co-expression of CD80 on tumor cells overcomes PD-L1 mediated suppression of T cell activity (Haile et al., 2011). Thus, in addition to its well-established function in triggering CD28 co-stimulation, CD80 upregulates T cell activity by neutralizing an inhibitory ligand in cis.

### PD-L1:CD80 *cis*-complex preserves the ability of CD80 to bind CD28

In the reciprocal experiment, we asked whether PD-L1:CD80 *cis*-interaction inhibits CD80:CD28 interaction. First, in a CD86 negative background, CD28-Fc stained CD80+PD-L1-Raji cells in a dose dependent manner (Figure 3A, red). Unexpectedly, co-expression of PD-L1 (5.9-fold higher than CD80, see Figure S4 for quantitation of expression levels) had no effect on the CD28-Fc binding at all concentrations tested (Figure 3A, blue), indicating that PD-L1 does not affect CD80:CD28 interaction. Consistent with this finding, in a T cell-SLB system in which CD80:CD28 *trans*-interaction is manifested by CD28 microclusters, addition of PD-L1 (three-fold excess to CD80) did not inhibit CD28 microcluster formation (Figure 3B). Furthermore, in the Raji-Jurkat co-culture assay, Raji (CD80+/CD86+/PD-L1-) and Raji (CD80+/CD86+/PD-L1+) cells led to indistinguishable degrees of CD28 and CD80 interfacial enrichment and CD28 phosphorylation (measured by co-immunoprecipitated p85) (Figure 3C, D). Importantly, PD-L1 was also enriched to the Raji-Jurkat interface in a CD80-dependent manner, suggesting that PD-L1, CD80, and CD28 form a tripartite complex that triggers CD28 signaling. Finally, knocking out CD80 from Raji significantly decreased the interfacial enrichment of both PD-L1 and CD28 (Figure 3C, bottom), as well as CD28-mediated p85 recruitment (Figure 3D). The residual p85 recruitment under this condition was due to CD86:CD28 interaction, because knocking out both CD80 and CD86 abolished this signal (Figure 3E). Collectively, these data demonstrate that PD-L1 does not affect CD80:CD28 *trans-*interaction and PD-L1:CD80 heterodimer is fully capable of activating CD28.

**Figure 3.**
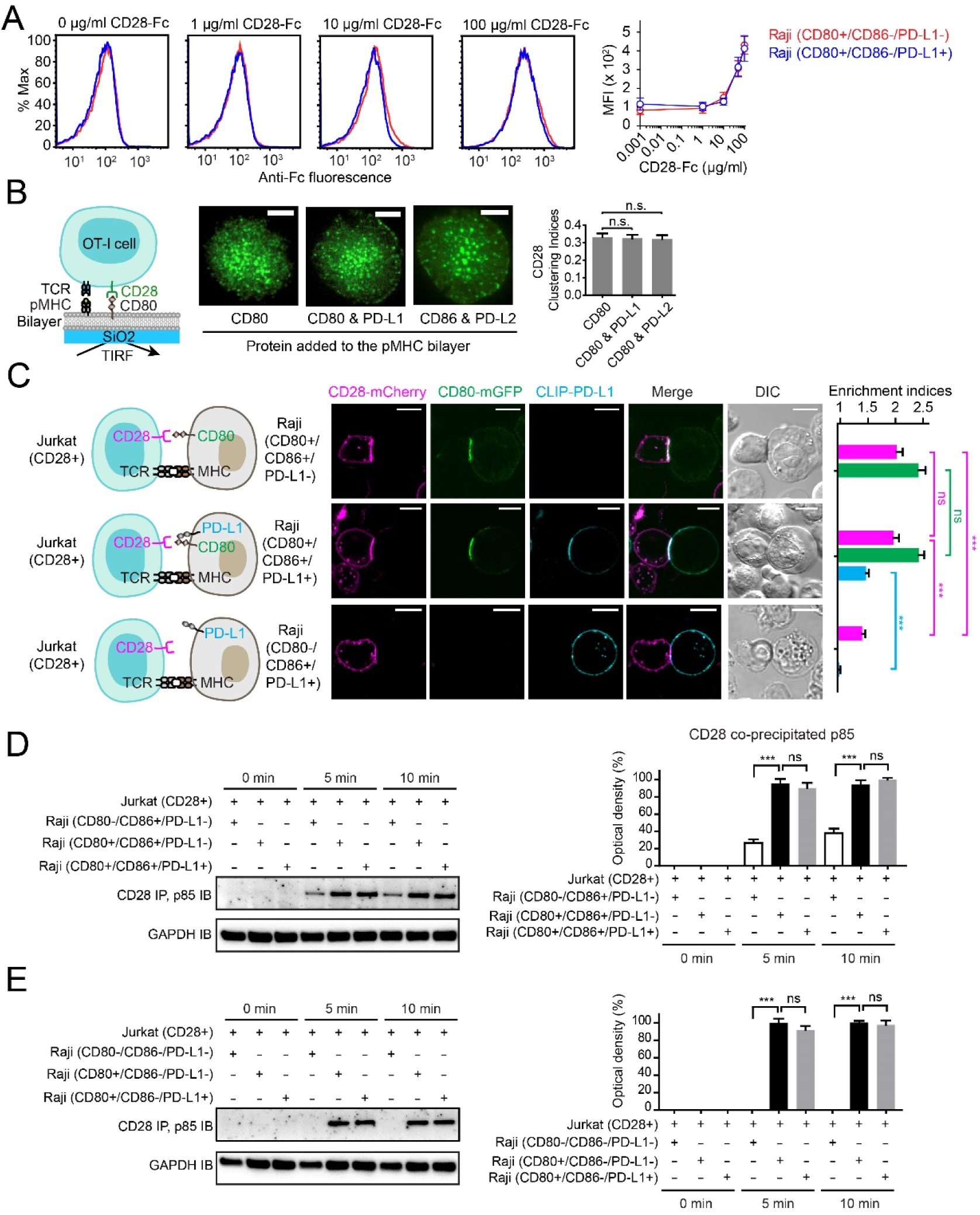
PD-L1:CD80 *cis*-interaction does not affect CD80:CD28 interaction. **(A)** Representative flow cytometry histograms of CD28-Fc staining to the indicated types of Raji B cells at the indicated concentrations. The MFI of bound CD28-Fc labeled by anti-human IgG AF647 was plotted against CD28-Fc concentration (means ± SEM, n = 4). **(B)** As depicted in the cartoon, CD28–mGFP transduced OT-1 cells interact with SLB functionalized with pMHC & CD80, pMHC & CD80 plus PD-L1 (three-fold excess to CD80), or pMHC & CD80 plus PD-L2 (three-fold excess to CD80). Shown are representative TIRF images of CD28–mGFP distribution 30 seconds after the cell-SLB contact. Bar graph shows the clustering index of CD28 under each condition. Data are means ± SEM of at least 18 cells from three independent experiments. Scale bars: 5 μm. See also Figure S2 for the corresponding images of TCR. **(C)** Cartoons on the left depicts CD28–mCherry transduced Jurkat conjugating with Raji expressing CD80–mGFP (CD80+/CD86+/PD-L1-), Raji expressing both CD80–mGFP and CLIP-PD-L1 (CD80+/CD86+/PD-L1+), or CD80 knock-out Raji expressing CLIP-PD-L1 (CD80-/CD86+/PD-L1+). Next to each cartoon are representative confocal images of the cell conjugate acquired two minutes after cell-cell contact, with CD28–mCherry shown in magenta, CD80–mGFP in green, and Dy647 labeled CLIP-PD-L1 in cyan. Scale bars: 10 µm. Bar graph on the right summarizes the synaptic enrichment indices (**Methods**) of the three conditions. Data are means ± SEM, n = 30 cells from three independent experiments. **(D)** Shown on the left are representative immunoblots showing CD28 co-immunoprecipitated (IP) p85 (PI3 kinase regulatory subunit) from the lysates of the indicated cocultures, with the times of lysis indicated (see **Methods**). WCL: whole cell lysate. Shown on the right is a quantification bar graph (means ± SEM, n = 3). **(E)** Same as **(D)** except CD86 was knocked out from all three types of Raji types. See also Figure S2 and Figure S4.

### PD-L1 protects CD80 from CTLA4 mediated *trans*-endocytosis

We next asked if the PD-L1:CD80 *cis*-interaction impacts the CD80:CTLA4 interaction. At a CD86 negative background, CTLA4-Fc stained CD80+PD-L1-Raji cells in a dose dependent manner (Figure 4A, red). Co-expression of PD-L1 (5.9-fold excess to CD80, see Figure S4 for quantitation of expression levels) substantially decreased CD80-mediated CTLA4-Fc staining (Figure 4A, blue), resulting in a 4.7-fold higher apparent dissociation constant (*K*_D_, 0.52 ± 0.16 μg/ml versus 0.11 ± 0.03 μg/ml). This result demonstrates that PD-L1:CD80 *cis*-interaction inhibits CD80:CTLA4 interaction, contrasting to the lack of effect on CD80:CD28 interaction. Our attempt to verify this finding in the T cell–SLB assay by probing CTLA4 plasma membrane microclusters was precluded by the intracellular localization of CTLA4, consistent with previous reports (Alegre et al., 1996; Linsley et al., 1996; Qureshi et al., 2012; Valk et al., 2008). Indeed, a major function of CTLA4 is to bind and *trans*-endocytose CD80 and CD86 (Hou et al., 2015), causing their depletion from APCs and weaker CD28 activation. Based on our finding that PD-L1 inhibits CD80:CTLA4 interaction, we predicted that PD-L1 protects CD80 from CTLA4-mediated depletion. To test this, we established a CTLA4-mediated *trans*-endocytosis assay: CTLA4–mGFP transduced Jurkat, but not wild-type Jurkat, decreased CD80 levels on Raji cells upon 30 minutes of cell contact (Figure 4B), indicating that CTLA4 *trans*-endocytosed CD80 from Raji APCs. Remarkably, co-expression of PD-L1 on Raji APCs significantly restored the CD80 level (Figure 4B). Thus, PD-L1 protects CD80 from CTLA4-mediated depletion.

**Figure 4.**
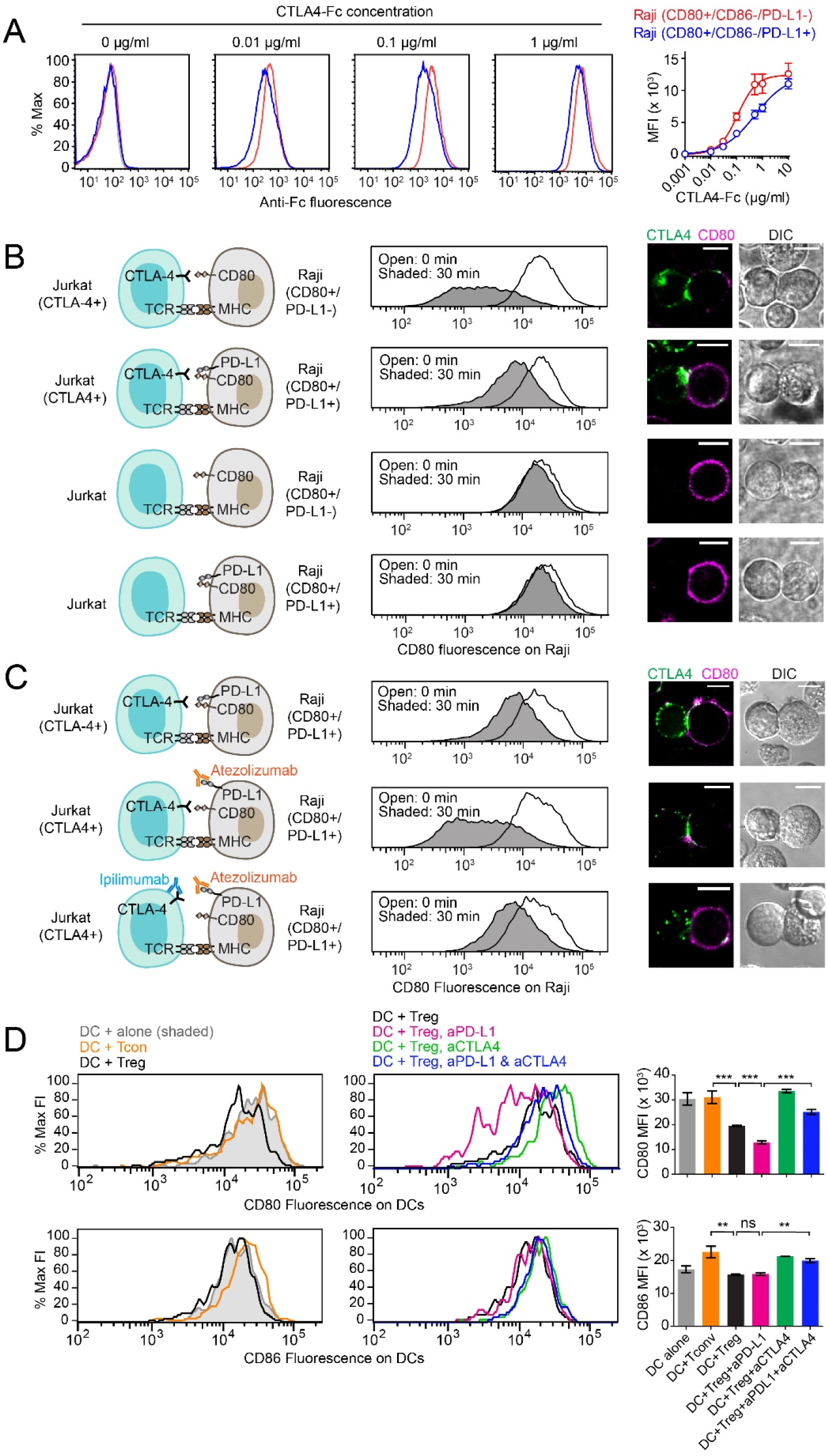
PD-L1:CD80 *cis*-interaction inhibits CD80:CTLA4 interaction and protects CD80 from CTLA4 mediated depletion. **(A)** Representative flow cytometry histograms of CTLA4-Fc binding to the three indicated types of Raji B cells at the indicated concentrations. The MFI of bound CTLA4-Fc labeled by anti-human IgG AF647 was plotted against [Ctla4-Fc] (means ± SEM, n = 3), and fit with “one site – specific binding” model using Graphpad Prism to obtain *K*_D_ values. **(B)** A Jurkat–Raji co-culture assay analyzing how PD-L1 interferes with CTLA4 mediated CD80 depletion. Shown on the left are cartoons depicting the co-cultured cells: CTLA4–mGFP transduced Jurkat (CTLA4+) or wild-type Jurkat (CTLA4-) were co-cultured with either PD-L1–mCherry transduced Raji (CD80+/PD-L1+) or wild-type Raji lacking PD-L1(CD80+/PD-L1-) (see **Methods**). Shown in the middle are flow cytometry histograms of CD80 expression (anti-CD80 APC) on Raji cells before (0 min) and after co-culture (30 min) and on the right are representative confocal images for the Jurkat–Raji conjugate from three independent replicates. Scale bars:10 μm. **(C)** An independent Jurkat–Raji conjugation assay examining how PD-L1 and CTLA4 blocking antibodies affect CD80 levels. Experiments were conducted as in **(B)** except pretreating the indicated cell type with atezolizumab (blocking PD-L1:CD80 *cis*-interaction) or ipilimumab (blocking CTLA4:CD80 *trans*-interaction). Shown in the middle and right are representative flow cytometry histograms of CD80 expression (anti-CD80 APC) and confocal images for the Jurkat–Raji conjugate from three independent replicates. Scale bars: 10 μm. **(D)** Flow cytometry histograms of CD80 and CD86 surface expressions on splenic DCs co-cultured with Treg cells in the presence or absence of a PD-L1 blockade antibody and/or a CTLA4 blockade antibody for 16 hr. Shown are representative flow cytometry histograms for CD80 and CD86 levels on DCs and bar graphs summarizing the result from three independent replicates. See also Figure S4.

### PD-L1 blockade antibodies downregulate CD80 on APCs in a CTLA4 dependent manner

A key prediction from our finding is that the FDA-approved PD-L1 blockade antibody atezolizumab, which blocks PD-L1:CD80 *cis*-interaction (Figure 1D), would deplete CD80 from APCs in a CTLA4-dependent manner. Consistent with notion, we found that atezolizumab treatment decreases CD80 on Raji APCs, and strikingly, this effect was substantially reversed by the CTLA4 blockade antibody ipilimumab (Figure 4C). We further test this model in a coculture system containing primary mouse dendritic cells (DC) and regulatory T cells (Treg), a type of suppressive T cells that downregulate CD80 and CD86 in *vivo* through CTLA4-mediated *trans*-endocytosis (Hou et al., 2015; Schmidt et al., 2009). Indeed, Tregs, but not conventional T cells (Tcon), decreased CD80 levels on DCs. The presence of a PD-L1 blockade antibody further decreased the CD80 level, and the co-administration of a CTLA4 blockade antibody restored the CD80 level. By contrast, CD86 level was unaffected by anti-PD-L1 treatment (Figure 4D). Finally, treatment of CT26 colon carcinoma implanted BALB/c mice with a PD-L1 blockade antibody led to a significant decrease of CD80, but not CD86 levels on tumor-infiltrating DCs and macrophages (Figure 5). Notably, administration of the PD-1 blockade antibody did not affect CD80 or CD86 levels on these cells (Figure 5). This finding illustrate an important difference between PD-L1 and PD-1 blockade antibodies. Collectively, data presented in this section uncover a role for PD-L1 in protecting CD80 from CTLA4-mediated *trans*-endocytosis and a CD80 depletion effect of PD-L1 blockade antibodies.

**Figure 5.**
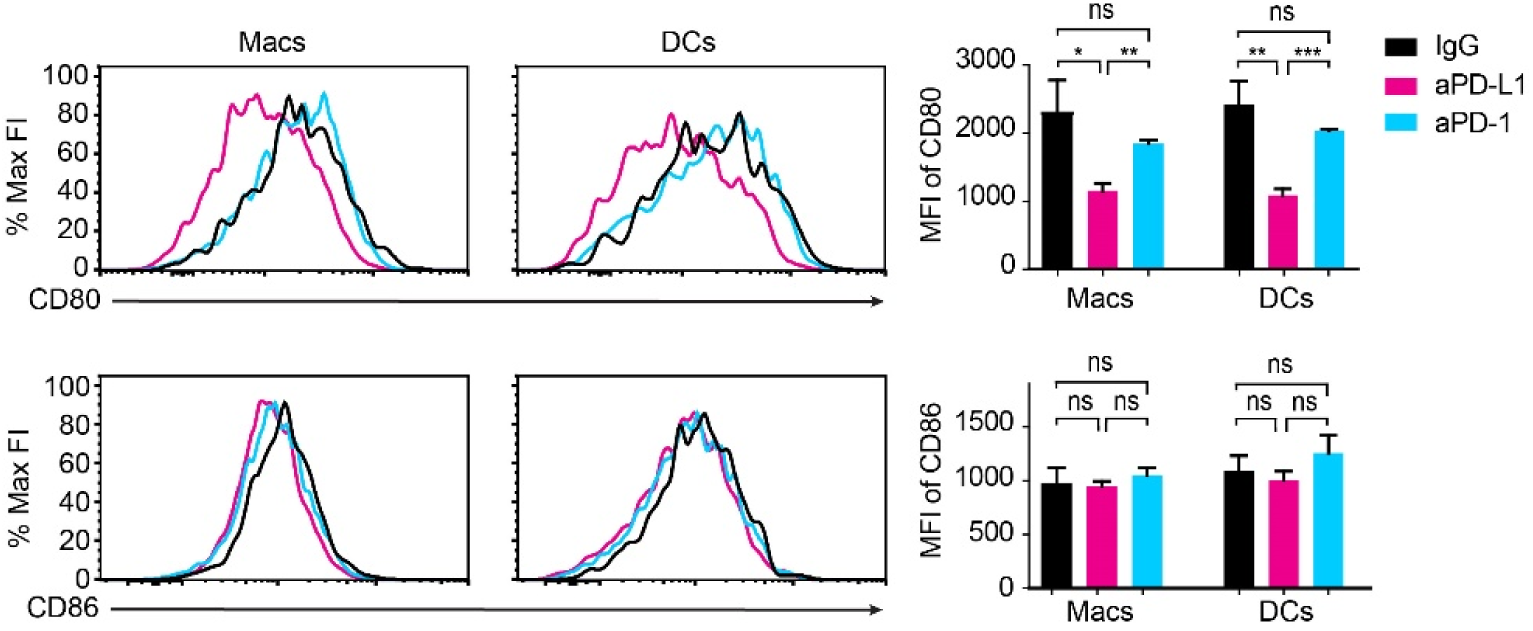
Anti-PD-L1, but not anti-PD-1 downregulates CD80 on tumor-infiltrating APCs. Shown on the left are flow cytometry histograms of CD80 and CD86 expressions on DCs and macrophages (Macs) isolated from tumor tissues of CT26-implanted BALB/C female mice treated with either PD-L1 blockade antibody (anti-PD-L1), PD-1 blockade antibody (anti-PD-1), or control IgG (see Methods). Shown on the right are bar graphs summarizing the MFI of CD80 and CD86 staining under the indicated condition.

## DISCUSSION

More than a decade after the first documentation of PD-L1:CD80 interaction (Butte et al., 2007), the biochemical and functional consequence of this interaction is still a mystery. The ability of CD80-expressing cells to bind PD-L1 coated surface led Butte et al. and the field to assume a *trans*-interaction model. By contrast, Freeman and colleagues recently proposed a *cis*-nature of this interaction (Chaudhri et al., 2018). In addition, the number of molecules involved (PD-1, CTLA-4, CD28, PD-L1, CD80) and the expression of PD-L1 and CD80 on both APCs and T cells has made it extremely challenging to determine the roles of PD-L1:CD80 interaction.

Using in *vitro* and cellular reconstitution assays, we demonstrate that PD-L1 and CD80 bind in *cis* but not in *trans*, thereby substantially simplifying the PD-L1:CD80:PD-1:CD28:CTLA4 network. We then discover that PD-L1:CD80 *cis*-interaction on APCs simultaneously represses two major immune checkpoints: PD-L1:PD-1 and CD80:CTLA4 pathways, while leaving CD80:CD28 interaction unaffected.

Conceivably, a substantial population of PD-L1 on APCs exist as *cis*-heterodimers with CD80, and *vice versa*. PD-L1:CD80 heterodimers are unable to engage either PD-1 or CTLA4 (Figure 2 & Figure 4), but bind and activate CD28 equally well as free CD80 molecules (Figure 3, see proposed model in Figure 6). It is likely that CD80:CTLA4 interface but not CD80:CD28 interface overlaps with CD80:PD-L1 interface. Despite the lack of a crystal structure of CD80:CD28 complex, crosslinking and mass spectrometry analyses indicate that CD80:CD28 interface is structurally distinct from CD80:CTLA4 interface, with the prior driven by electrostatic interactions and the latter driven by hydrophobic interactions (Sorensen et al., 2004).

**Figure 6.**
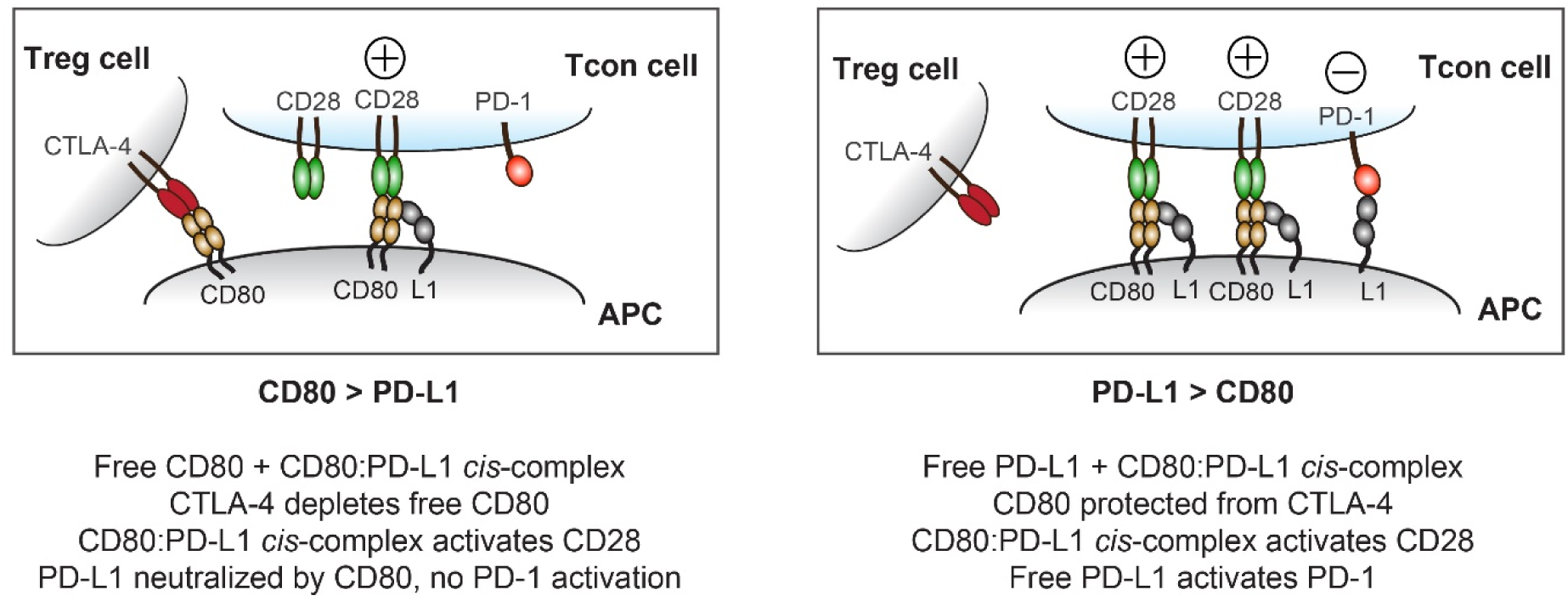
A proposal model for how CD80:PD-L1 heterodimerization regulates T cell signaling. Due to *cis*-interaction (Figure 1), CD80 and PD-L1 on the APCs can exist as three forms: free CD80, free PD-L1, and CD80:PD-L1 heterodimers. (*Left*) If an APC expresses higher CD80 than PD-L1, then CD80:PD-L1 heterodimers and free CD80 can be found, while free PD-L1 would be sparse. As the APC encounters a Treg cell, the free CD80 would be *trans*-endocytosed and depleted by Treg-specific CTLA-4. The PD-L1-bound CD80, however, is protected from CTLA-4 mediated depletion (Figure 4). These PD-L1-bound CD80 would instead activate CD28 on Tcon cells (Figure 3). Because CD80 blocks PD-L1:PD-1 interaction, CD80:PD-L1 complex cannot bind PD-1 either (Figure 2). (*Right*) If an APC surface has higher PD-L1 than CD80, then the two dominant forms would be PD-L1:CD80 heterodimers and free PD-L1, while free CD80 would be sparse. The PD-L1:CD80 heterodimer, which cannot be depleted by Treg-specific CTLA4 (Figure 4), would bind and activate CD28 but not PD-1 (Figure 2 & Figure 3). However, the free PD-L1 binds and activates PD-1 on T cells.

Our results uncover an unexpected role of PD-L1 in repressing the CTLA4 pathway. Independent to its well-established role in triggering PD-1 in *trans*, PD-L1 can block CD80:CTLA4 interaction through binding to CD80 in *cis*. Importantly, this allows PD-L1 to protect CD80 from CTLA4-mediated *trans*-endocytosis, thereby sustaining CD80 levels on APCs. From a different perspective, PD-L1 steers CD80 from CTLA4 to CD28, while free CD80 strongly prefers CTLA4 over CD28, the PD-L1 bound CD80 interacts with CD28 only (Figure 6). Though atezolizumab is best known to block PD-L1:PD-1 interaction, we show that it also block PD-L1:CD80 *cis*-interaction (Figure 1D). Consequently, atezolizumab frees up CD80 for CTLA4-mediated *trans*-endocytosis, causing CD80 downregulation on APCs. Thus, our study reveals two opposing effects of atezolizumab on T cell signaling: first, atezolizumab blocks PD-L1:PD-1 interaction to stimulate T cell activation; second, by dissolving PD-L1:CD80 heterodimers, atezolizumab switches CD80 from a CD28 ligand to a CTLA4 ligand, thereby inhibiting T cell activation. We speculate that the net outcome of atezolizumab depends on the relative expression levels of PD-1 and CTLA4. Our model predicts that as PD-1 levels increase on T cells, PD-L1 would switch from a positive regulator (CD80 protecting) to a negative regulator (PD-1 triggering). Importantly, despite the general assumption that anti-PD-1 and anti-PD-L1 therapies have identical clinical responses, we show that anti-PD-L1 but not anti-PD-1 depletes CD80 on APCs (Figure 5). Indeed, clinical trials have revealed that anti-PD-1 have a significantly higher overall response rate than anti-PD-L1 (Brahmer et al., 2012; Passiglia et al., 2018).

Data presented here provide a mechanistic rationale for co-blockade of PD-L1 and CTLA4 in cancer immunotherapy, a form of combination therapy currently under evaluation in clinical trials (Chae et al., 2018). As reported in Figure 4D, CTLA4 blockade negates the CD80-depleting effect of PD-L1 blockade. Our finding also suggests that PD-L1 inhibitors that selectively block PD-L1:PD-1 interaction, but not PD-L1:CD80 interaction, might prove more effective than atezolizumab and other PD-L1:PD-1/PD-L1:CD80 dual blockers (e.g., durvalumab, avelumab) in immunotherapy.

The present study also elucidates that by sequestering PD-L1 in *cis*, CD80 on APCs functions as a rheostat for PD-L1:PD-1 signaling (Figure 6), suggesting CD80 as a biomarker for PD-L1:PD-1 targeted immunotherapy. Currently, PD-L1 expressions on tumors and tumor infiltrating APCs are often used to predict patient response for PD-1 targeted therapy (Kluger et al., 2017; Lin et al., 2018; Tang et al., 2018). Despite a positive correlation between PD-L1 expression and patient response, many PD-L1 positive patients fail to respond to PD-1 or PD-L1 blockade antibodies. Our study suggests that patients with high CD80 expression on tumor infiltrating APCs would less likely respond to PD-L1 or PD-1 inhibitors even though they are PD-L1 positive, because CD80 would mask PD-L1 in *cis* to prevent PD-1 activation in the first place. It is also worth noting that CD80 can block at least some clones of PD-L1 staining antibodies, leading to a substantial underestimation of PD-L1 levels on CD80+ cells (Figure S1A). Therefore, choosing the right clones of PD-L1 staining antibodies would be essential for reliably probing PD-L1 levels in tumor tissues.

## METHODS

### Cell cultures

HEK293T cells and Raji B cells were obtained from Dr. Ronald Vale (University of California San Fran*cis*co), Jurkat T cells from Dr. Arthur Weiss (University of California San Fran*cis*co), HEK293F cells from Dr. Andrew Ward (Scripps Research). HEK293T cells were maintained in DMEM medium (Genesee Scientific, catalog 25-501) supplemented with 10% fetal bovine serum (Omega Scientific, catalog FB-02), 100 U/mL of Penicillin (GE Healthcare, catalog SV30010), and 100 µg/mL of Streptomycin (GE Healthcare, catalog SV30010)) at 37 °C/5% CO_2_. Jurkat T cells and Raji B cells were maintained in RPMI medium (corning, catalog 10-041-CM) supplemented with 10% fetal bovine serum, 100 U/mL of Penicillin, and 100 µg/mL of Streptomycin) at 37 °C/5% CO_2_. HEK293F cells were maintained in FreeStyle 293 Expression Medium (Thermo Fisher Scientific, catalog 12338018) at 37 °C/8% CO_2_. OT-1 splenocytes were harvested from C57BL/6-Tg (TcraTcrb) 1100Mjb/J (OT-1) mice (Jackson Laboratory) and maintained in OT-1 culture medium (RPMI 1640 supplemented with 10% fetal bovine serum, 1 mM Sodium Pyruvate (Corning, 25-000-CI), 50 µM β-mecaptoethanol (Fisher Scientific, catalog ICN19024283), 100 U/mL of Penicillin, and 100 µg/mL of Streptomycin) at 37 °C/5% CO_2_.

### Recombinant proteins

Recombinant His-tagged mouse and human PD-L1, PD-L2, CD80, CD86, and ICAM used in OT-1–SLB assay or LUVs-SLB adhesion assay were purchased from Sino Biological. For FRET assay using LUVs, extracellular portion (ectodomain) of human PD-L1 (aa 19–239), human PD-L2 (aa 20–220), or human CD80 (aa 35–242) was expressed in HEK293F cells, as described previously(Murin et al., 2014). The N terminus of each ectodomain was fused with the signal peptide of HIV envelope glycoprotein gp120 followed by a twinstrep tag (amino acids sequence: WSHPQFEKGGGSGGGSGGSAWSHPQFEK) and a SNAP-tag. The C terminus of each extracellular segment was fused with a decahistidine (His) tag. For His-tag free CD80, the His-tag coding sequence was removed from the expression construct. All His-tagged proteins were purified from the cell culture medium using HisTrap Excel column (GE Healthcare, catalog 17371206) and eluted using buffer containing 50 mM HEPES-NaOH, pH 8.0, 150 mM NaCl, 0.5 M imidazole. His10-tag free CD80 ectodomain was purified with a StrepTrap HP column (GE Healthcare, catalog 28907547) in 100 mM Tris-HCl, 150 mM NaCl, 1 mM EDTA, pH 8.0 and eluted with the same buffer containing 2.5 mM desthiobiotin (Sigma-Aldrich, catalog D1411). The ectodomain of mouse MHC-I molecule H2-Kb was produced as a disulfide-stabilized single chain trimer with a covalently linked ovalbumin (OVA) peptide SIINFEKL (Mitaksov et al., 2007), and a C-terminal His-tag, using the Bac-to-Bac baculovirus expression system, as previously described (Hui et al., 2017). All affinity-purified proteins were size-exclusion-purified using a Superdex 200 Increase 10/300 GL column (GE Healthcare, catalog 28990944) in HEPES buffered saline (50 mM HEPES-NaOH, pH 7.5, 150 mM NaCl, 10% glycerol). Gel filtered proteins were labeled with either SNAP-Cell 505 (NEB, catalog S9103S) or SNAP-Cell TMR (NEB, catalog S9105S) following manufacturer’s instructions. Free dyes were then removed using Zeba Spin Desalting Columns (Thermo Fisher Scientific, catalog P187769). All proteins were quantified by SDS-PAGE and Coomassie blue staining, using bovine serum albumin (BSA, Thermo Scientific, catalog 23209) as a standard.

### Cell lines generation

To generate CD80 KO Raji cells (CD80-/CD86+/PD-L1-), two PX330-GFP vectors coding different CD80 sgRNA were electroporated into wild-type Raji cells (CD80+/CD86+/PD-L1-) using Cell Line Nucleofector Kit V (LONZA, catalog VACA-1003). Electroporated cells were recovered for two days at 37 °C/5% CO_2_. GFP-positive cells were then sorted by flow cytometry and maintained in culture media for one week, after which the cells were stained with anti-CD80 APC (Biolegend, catalog 305220) and CD80 KO cells were sorted by flow cytometry. CD86 KO Raji cells (CD80+/CD86-/PD-L1-) were generated in the same manner as CD80. CD80/CD86 double KO Raji cells (CD80-/CD86-/PD-L1-) were generated by knocking out CD80 from CD86 KO Raji cells. Cells were sorted with anti-CD80 APC and anti-CD86 BV421 (Biolegend, catalog 305425) staining.

Each gene of interest was introduced into Jurkat and Raji cells via lentiviral transduction, as described previously (Zhao et al., 2018). For Figure 2 and Figure S1, Raji (PD-L1+/CD80 KO) and Raji (PD-L1+/CD80 low) were generated by transducing PD-L1–mCherry to CD80 KO Raji (CD80-/CD86+/PD-L1-) and wild-type Raji (CD80+/CD86+/PD-L1-), respectively. PD-L1–mCherry transduced wild-type Raji cells were further transduced with CD80–CLIP to produce CD80-overexpressing Raji (Raji (PD-L1+/CD80 high)). Jurkat (PD-1+) was generated previously by transducing PD-1– mGFP into Jurkat wild-type (Hui et al., 2017). For Figure 3A, 3E, 4A and Figure S4, Raji (CD80+/CD86-/PD-L1+) cells were generated by transducing PD-L1–mCherry into CD86 KO Raji cells (CD80+/CD86-/PD-L1-). For Figure 3C, Jurkat (CD28+) cells were generated by transducing Jurkat wild-type with CD28–mCherry. Raji (CD80+/CD86+/PD-L1-), Raji (CD80+/CD86+/PD-L1+), and Raji (CD80-/CD86+/PD-L1+) were generated by transducing dSV40-promoter-driven CD80–mGFP and/or CLIP-PD-L1 into CD80 KO Raji (CD80-/CD86+/PD-L1-) cells. For Figure 3D, Raji (CD80+/CD86+/PD-L1+) were the same as Raji (PD-L1+/CD80 low) used in Figure 2 that express low, endogenous levels of both CD80 and CD86. Wild-type Raji (CD80+/CD86+/PD-L1-) and CD80 KO Raji (CD80-/CD86+/PD-L1-) were used in the same experiment. For *trans*-endocytosis assay in Figure 4B, C, Jurkat (CTLA4+) was generated by introducing CTLA4–mGFP to Jurkat wild-type via lentivirus under the control of the dSV40 promoter. Wild-type Raji (CD80+/PD-L1-) and PD-L1–mCherry-transduced Raji (CD80+/PD-L1+) were used for *trans*-endocytosis assay.

### Confocal microscopy based FRET assay with HEK293T cells

pHR plasmid encoding full length human PD-L1 or PD-L2 with an N-terminally fused CLIP tag (CLIP-PD-L1, CLIP-PD-L2) was co-transfected with pHR encoding either SNAP-tagged full length CD80 (SNAP-CD80) or CD86 (SNAP-CD86) into HEK293T cells using polyethylenimine (Fisher Scientific, catalog NC1014320). 72-hour post transfection, cells were trypsinized and seeded on Poly-D-lysine (Sigma-Aldrich, catalog P6407) treated 96-wells plate with a glass bottom (Dot Scientific, catalog MGB096-1-2-LG-L). 24 hours later, cells were labeled with CLIP-Surface 547 (NEB, catalog S9233S) and SNAP-Surface Alexa Fluor 647 (NEB, catalog S9136S) at 37 °C/ 5% CO_2_ for 30 minutes, and washed 3 times with 1× PBS (pH 7.4). Labeled cells were then fixed with 4% paraformaldehyde (PFA, Fisher Scientific, catalog 50980494) and used for the FRET assay. Images were acquired with an Olympus FV1000 confocal microscope by exciting CLIP-Surface 547 (energy donor) at 543 nm and SNAP-Surface Alexa Fluor 647 (energy acceptor) at 635 nm. Donor images before and after acceptor photobleaching were acquired for FRET analysis using ImageJ (Fiji) with the AccPbFRET plugin, as described (Roszik et al., 2008).

### FRET assay with protein reconstituted LUVs

Synthetic 1,2-dioleyl-sn-glycero-3-phosphocholine (POPC, catalog 850457C) and 1,2-dioleoyl-sn-glycero-3-[(N-(5-amino-1-carboxypentyl) iminodiacetic acid) succinyl] (nickel salt, DGS-NTA-Ni, catalog 790404C) were purchased from Avanti Polar Lipids. LUVs composing of 80% POPC and 20% DGS-NTA-Ni were generated by the extrusion method, as described(Hui and Vale, 2014). Briefly, desired lipids were mixed in chloroform, dried under a nitrogen stream and desiccated in a vacuum container for one hour. To form LUVs, the dried lipid film was resuspended with PBS and extruded for 20 times through a pair of polycarbonate filters containing pores of 200 nm diameter. For Figure 1D, 0.23 nM LUVs in PBS containing 1.5 mg/mL BSA and 1 mM TCEP were incubated at room temperature with 25 nM SC505-labeled PD-L1-His alone, with 25 nM SC505*PD-L1-His and 75 nM atezolizumab combined, or with 25 nM SC505*PD-L2-His alone, in a 96-well solid white microplate (Greiner Bio-One, catalog 655075), during which the SC505 fluorescence was monitored in real time using a plate reader (Tecan Spark 20) with 504-nm excitation and 540-nm emission. Following 90 minutes’ incubation, TMR-labeled CD80-His (TMR*CD80-His) or TMR*CD80 without His-tag was injected and SC505 fluorescence monitored for an additional 60 minutes. The average of last ten fluorescence value before the addition of TMR*CD80-His or TMR*CD80 (no His) was used to normalize the data.

### LUVs-SLB adhesion assay

To form SLB, a glass bottom 96-well plate was incubated with 5% Hellmanex III (Hëlma Analytics, catalog Z805939) overnight on a 50 °C heat pad, thoroughly rinsed with ddH_2_O and sealed with Nunc sealing tape (Thermo Fisher Scientific, catalog 232698). The desired wells were washed twice with 5 M NaOH (30 minutes each), and three times with 500 µL ddH_2_O followed by equilibration with PBS. Small unilamellar vesicles (SUVs, composing of 97.5% POPC, 2% DGS-NTA-Ni and 0.5% PEG5000 PE (1,2-dioleoyl-sn-glycero-3-phosphoethanolamine-N-[methoxy(polyethylene-glycol)-5000], ammonium salt (Avanti Polar Lipids, catalog 880230C)) were prepared as described previously (Hui et al., 2017) and added to the cleaned wells containing 200 µL 1× PBS, and incubated for 90 minutes at 50 °C and 30 minutes at 37 °C to induce SLB formation. The SLBs were then rinsed thoroughly with PBS to remove excess SUVs, and blocked with 1 mg/mL BSA in PBS for 30 minutes at 37 °C. The SLBs were then overlaid with 200 µL of either 3 nM human PD-1-His (Sino Biological, catalog 10377-H08H), 3 nM human CD80-His (Sino Biological, catalog 10698-H08H), or 3 nM human CD86-His (Sino Biological, catalog 10699-H08H). After 1-hour incubation at 37 °C, the unbound proteins were washed away using excess PBS containing 1 mg/mL BSA. The plate was incubated at 37 °C for another 30 minutes and washed again with PBS containing 1 mg/mL BSA to remove dissociated proteins, leaving the bilayer with stably bound proteins (Nye and Groves, 2008). LUVs containing both DGS-NTA-Ni and Bodipy-PE (89.7% POPC + 10% DGS-NTA-Ni + 0.3% Bodipy-PE (N-(4,4-Difluoro-5,7-Dimethyl-4-Bora-3a,4a-Diaza-s-Indacene-3-Propionyl)-1,2-Dihexadecanoyl-sn-Glycero-3-Phosphoethanolamine, Triethylammonium Salt (Thermo Fisher Scientific, catalog D3800)) were prepared by the aforementioned extrusion method. 0.23 nM LUVs with Bodipy-PE were then incubated with 8.3 nM human PD-L1-His (Sino Biological, catalog 10084-H08H) for 90 minutes in PBS with 1 mg/mL BSA at room temperature, to ensure stable, 100% protein binding. The protein-bound LUVs were then added onto either PD-1, CD80, or CD86 functionalized SLBs. After 10-minute incubation, unbound LUVs were washed away with excess PBS and the SLB-captured LUVs visualized and recorded by a Nikon Eclipse Ti TIRF microscope equipped with a 100× Apo TIRF 1.49 NA objective, controlled by the Micro-Manager software (Edelstein et al., 2014). The fluorescence intensity of LUVs from the Bodipy (488 nm) channel in the TIRF field was calculated using the ImageJ software.

### OT-1–SLB microscopy assay

OT-1 primary T cells were retrovirally transduced with either mouse PD-1–mCherry or mouse CD28–mGFP. Retroviruses were produced as described previously (Hui et al., 2017). Freshly harvested OT-1 splenocytes were stimulated with 10 nM SIINFEKL peptide (Anaspec, catalog AS-60193-1) in OT-1 culture medium supplemented with 100 U/ml mouse recombinant IL-2 (Thermo Fisher Scientific, catalog 14802164) in a 37 °C/5% CO_2_ incubator. 36 hours later, cells were resuspended in retrovirus supernatants containing 8 µg/ml Lipofectamine and 100 U/ml mouse recombinant IL-2, spin-infected at 35 °C, 1000 × g for 120 minutes, and incubated at 37 °C/5% CO_2_ overnight. The virus supernatant was replaced with fresh OT-1 culture medium supplemented with 10 nM SIINFEKL peptide and 100 U/ml mouse recombinant IL-2 the second day and cells incubated for another 48–96 hours before microscopy. For TIRF microscopy, the 96-well plate was treated with SUVs to form SLB as described above. For Figure 2B, SLB was functionalized by a mixture of 5 nM pMHC-I-His, 2 nM mouse ICAM-His (Sino Biological, catalog 50440-M08H) and either 3 nM mouse PD-L1-His (Sino Biological, catalog 50010-M08H), 3 nM mouse PD-L1-His/9 nM mouse CD80-His (Sino Biological, catalog 50446-M08H) combined, or 3 nM mouse PD-L1-His/9 nM mouse CD86-His (Sino Biological, catalog 50068-M08H) combined. For Figure 3B, SLB was functionalized by a mixture of 5 nM pMHC-I-His and 2 nM mouse ICAM-His and either 3 nM mouse CD80-His, 3 nM mouse CD80-His/9 nM mouse PD-L1-His combined, or 3 nM mouse CD80-His/9 nM mouse PD-L2-His (Sino Biological, catalog 50804-M08H) combined. Transduced OT-1 cells were harvested via centrifugation at 200 × g for 4 minutes, incubated with 10 µg/ml AF647-labeled mouse TCRβ antibody (Biolegend, catalog H57-597) for 30 minutes on ice, washed three times with imaging buffer (Hui et al., 2017), and then plated onto functionalized SLBs. TIRF images were acquired at 37 °C on a Nikon Eclipse Ti microscope equipped with a 100× Apo TIRF 1.49 NA objective, controlled by the Micro-Manager software and analyzed with ImageJ. Clustering index was calculated by dividing the fluorescence intensity of PD-1 or CD28 microclusters by the total fluorescence intensity of the respective channel of the entire cell.

### Jurkat–Raji conjugation assay

For cell conjugation assay, Raji B cells were pre-incubated with 30 ng/mL SEE superantigen alone (Figure 2C, Toxin Technology, catalog ET404) or SEE together with 1 µM CLIP-Surface 647 (Figure 3C, NEB, catalog S9234S) in RPMI medium for 30 minutes at 37 °C. Cells were then washed twice to remove free SEE and dye. Following antigen loading, 0.4 million antigen-loaded Raji B cells and 0.4 million Jurkat T cells were precooled on ice and mixed in a 96-well plate. The plate was then centrifuged at 290× g for one minute at 4 °C to initiate cell–cell contact, and immediately transferred to a 37 °C water bath. Two minutes later, cells were resuspended and fixed with 1% PFA and loaded into a 96-well glass-bottom plates for confocal microscopy assays. Images were acquired with an Olympus FV1000 confocal microscope and processed, and quantified using ImageJ. Interface enrichment indices of PD-1 and CD28 on Jurkat cells and PD-L1 and CD80 on Raji cells were computed by dividing the fluorescence density at the interface divided with fluorescence density of the cell membrane excluding the interface. Fluorescence density was calculated as fluorescence intensity divided by area. The interface was defined as the conjugated area between Jurkat and Raji cells based on the DIC images.

For examining receptor phosphorylation in Figure 2D, Figure 3D, E. Serum starved Jurkat cells and SEE-loaded Raji cells (2 million each) were co-pelleted as described above. The cell pellets were lysed with ice cold NP40 buffer (50 mM HEPES, pH 7.5, 300 mM NaCl, 2% NP40, 2 mM EDTA, 10% glycerol, 2 mM PMSF, 20 mM Na_3_VO_4_, 20 mM NaF) at indicated time points. For 0 minute samples, Raji-Jurkat mixtures were lysed prior to centrifugation. Lysates were centrifuged at 15, 000x g for 10 minutes. PD-1–mGFP or CD28 were immunoprecipitated by using GFP-Trap (Chromotek, catalog gta-20) and anti-CD28 antibody (Bio X Cell, catalog BE0291) coated Protein G Dynabeads (Thermo Fisher Scientific, catalog 10004D), respectively. Equal fractions of the immunoprecipitates were subjected to SDS-PAGE and blotted with anti-p85 antibody (Cell Signaling Technology, catalog 4292), anti-phosphotyrosine antibody (Sigma-Aldrich, catalog P4110), or GFP antibody (Thermo Fisher Scientific, catalog A6455). Whole cell lysates (WCL) were blotted with anti-GAPDH antibody (Proteintech, catalog 10494-1-AP).

### Raji B cell staining with Fc-fusion proteins

Raji B cells were incubated with either PD-1-Fc, CD28-Fc, or CTLA4-Fc at indicated concentrations (Figure 2A, 3A, and 4A) for 35 minutes on ice, washed twice with PBS plus 2% FBS, and stained with an AF647-labeled anti-human IgG Fc antibody (Biolegend, catalog 409320). Cells were then analyzed by flow cytometry on LSRFortessa analyzer (BD Biosciences). The data were analyzed by FlowJo and plotted by GraphPad Prism 5.

### Quantification of PD-L1 and CD80 expression

For flow cytometry based quantification, PD-L1 and CD80 were stained by anti-PD-L1 PE (eBioscience, catalog 14-5983-82) and anti-CD80 PE (Biolegend, catalog 305208) and their expression levels were quantified using the QUANTUM™ R-PE MESF kit (Bangs Laboratories Inc, catalog 827), following manufacturer’s instructions. For western blot-based quantification, total cell lysate was applied for SDS-PAGE and detected by anti-PD-L1 PE (eBioscience, catalog 14-5983-82) and anti-CD80 (Novus Biologicals, NBP2-25255). The latter was further detected by using DyLight488 anti-mouse IgG (Biolegend, catalog 405310). Molecular densities were calculated assuming the following diameters:13 μm for Raji B cells (Hui et al., 2017) and 12.5 μm for DCs (Dumortier et al., 2005).

### Flow cytometry analysis of human monocyte-derived DCs

Human peripheral blood CD14+ monocytes were isolated from normal human peripheral blood (iXCells, catalog 10HU-008), and cultured in RPMI medium supplemented with 50 ng/ml of GM-CSF and 50 ng/ml of IL-4 in 37 °C/5% CO_2_. After five-day incubation, when the majority of monocytes were differentiated to immature DCs, cells were re-cultured in RPMI medium supplemented with 50 ng/ml TNF-α for an additional two days to generate mature DCs. PD-L1 and CD80 expression levels on both immature and mature DCs were measured by flow cytometry. Briefly, cells were pre-incubated with Human TruStain FcX™ (BioLegend, catalog 422301) to block Fc receptors, and then incubated with the viability dye Ghost Dye Violet 450 (Tonbo Biosciences, catalog 10140-978), followed by an antibody mixture containing anti-CD1a PerCP/Cy5.5 (Biolegend, catalog 300129), anti-CD14 FITC (Biolegend, catalog 301804), and anti-PD-L1 PE (eBioscience, catalog 14-5983-82) or anti-CD80 PE (Biolegend, catalog 305208). Stained cells were processed on a BD LSRFortessa cell analyzer and flow cytometry data were analyzed by FlowJo software.

### *Trans*-endocytosis assay

For analyzing CD80 *trans*-endocytosis, CTLA4–mGFP expressing Jurkat or wild-type Jurkat cells were co-cultured with either PD-L1–mCherry expressing Raji or wild-type Raji. Briefly, 0.4 million SEE-loaded Raji B cells and 0.4 million Jurkat T cells were mixed and co-pelleted by centrifugation at 290× g for one minute, and incubated at 37°C/5% CO_2_ for 30 minutes. After incubation, cells were resuspended and stained with anti-CD80 APC, anti-CD3 PE (Biolegend, 317308), and anti-CD20 PE/Cy7 (Biolegend, catalog 302311) for flow cytometry with a BD LSRFortessa cell analyzer. CD3 and CD20 were used to gate Raji cells out from cell mixture. CD80 expression level was then analyzed on Raji. For blockade treatment in Figure 4C, both SEE-loaded Raji and Jurkat were treated with 2 μg of either atezolizumab (Selleckchem, catalog A2004) or ipilimumab (Selleckchem, catalog A2001) per million cells for 15 minutes prior to mixing, and blockade antibodies were kept in the coculture until the staining step. For confocal microscopy, stained cells were plated on Poly-D-lysine treated glass bottom 96-well plate and images were acquired with an FV1000 confocal microscope in GFP and APC channels.

### DCs-Treg cells co-culture Assay

Tcon cells and Treg cells were isolated using magnetic bead isolation system. In brief, total CD4+ T cells were first isolated from Foxp3^Thy1.1^ reporter mice(Liston et al., 2008) by using mouse CD4 T cell isolation kit (Biolegend, catalog 48006), and stained with anti-Thy1.1-PE antibody (eBioscience, catalog 12-0900-83). Next, Treg cells were further separated with Tcon cells by using anti-PE beads (Miltenyi Biotec, catalog 130-048-801). DCs were isolated from Ly5.1+ mice by using CD11c isolation kit (Miltenyi Biotec, catalog 130-108-338). DCs (5 x 10^3^) were cultured with either Tcon cells (5 x 10^4^) or Treg cells (5 x 10^4^) in the presence or absence of anti-PD-L1 (50 µg/ml, Bio X Cell, catalog BE0101) and/or anti-CTLA4 (50 µg/ml, Bio X Cell catalog BE0164) in RPMI 1640 supplemented with 10% fetal bovine serum, 50 µM β-mecaptoethanol, 100 U/mL Penicillin, and 100 µg/mL Streptomycin, 0.5 µg/ml Anti-CD3 mAb (Bio X Cell, catalog BE0001-2) and 1 µg/ml LPS (Enzo Life Sciences, catalog ALX-581-008-L002). After 16 hours, cells were first stained with Ghost Dye Red 780 (Tonbo Biosciences, catalog 13-0865-T100), followed by surface antibody staining for Ly5.1 (eBioscience, catalog 11-0453-82); Thy1.1 (eBioscience, catalog 12-0900-83); CD11c (eBioscience, catalog 45-0114-82); CD80 (Biolegend, catalog 104733); CD86 (eBioscience, catalog 17-0862-82); and PD-L1 (Biolegend, catalog 124315). An LSRFortessa analyzer (BD Biosciences) was used for data collection, and FlowJo software was used for data analysis.

### Anti-PD-L1 treatment of CT26 tumor bearing mice

8-12 weeks old BALB/C female mice were transplanted with 2 × 10^6^ CT26 cells in the subcutaneous space. 400 μg of anti-PD-1 (Bio X Cell, catalog BE0273), anti-PD-L1 (Bio X Cell, catalog BE0101), or control IgG (Bio X Cell, catalog BE0090) were intraperitoneally injected five days later. 24 hours after antibody injection, the tumors were harvested from all animals, homogenized, and stained for expression of CD80 or CD86 on live (7AAD-) dendritic cells (CD11b+ CD11c+) or macrophages (CD11b+ F4/80+). All antibodies used for flow cytometry were purchased from Biolegend, including anti-CD11b (Biolegend, catalog 101225). anti-CD11c (Biolegend, catalog 117317), anti-F4/80 (Biolegend, catalog 123113), anti-CD80 (Biolegend, catalog 104713), anti-CD86 (Biolegend, catalog 105005), and anti-PD-L1 (Biolegend, catalog 124307). An FACSCantoII analyzer (BD Biosciences) was used for data collection. FlowJo software was used for data analysis.

### Quantification and statistical analysis

Unless otherwise indicated, data were reported as mean ± SEM, and number of replicates were indicated in figure legends. Curve fitting and normalization were performed in GraphPad Prism 5. Statistical significance was evaluated by two-tailed *Student’s* t test (*, p < 0.05; **, p < 0.01; ***, p < 0.001) in GraphPad Prism 5. Data with p < 0.05 are considered statistically significant.

## ACKNOWLEDGEMENTS

We thank E. Bennett (UCSD) for critically reading the manuscript; C. Bonorino for help in deriving DCs from monocytes, N. Stuurman for technical support with TIRF microscopy; J. Sabatini and M. Bohm for help with confocal microscopy at the UCSD School of Medicine Microscopy Core supported by NINDS P30 Grant (NS047101). Research reported here was supported by National Institute of Health, National Cancer Institute, under award number R37CA239072. Y. Zhao is a Cancer Research Institute Irvington Postdoctoral Fellow. E. Hui is a Searle Scholar and a Pew Biomedical Scholar.

## AUTHOR CONTRIBUTIONS

Y.Z. and E.H. designed the project. C.H.L. performed the DC-Treg coculture assay. C.L. and J.B. conducted the experiments related to CT26. X.X. performed experiments for data in Figure 2D, Figure 3D and Figure 3E. Z.H. assisted experiments in Figure 2B and Figure 3B. Y.Z. conducted all the other experiments and data analyses. C.X., J.B., L.F.L. and E.H. supervised the project. Y.Z. and E.H. prepared the manuscript with input from all authors.

---

**Figure S1.**
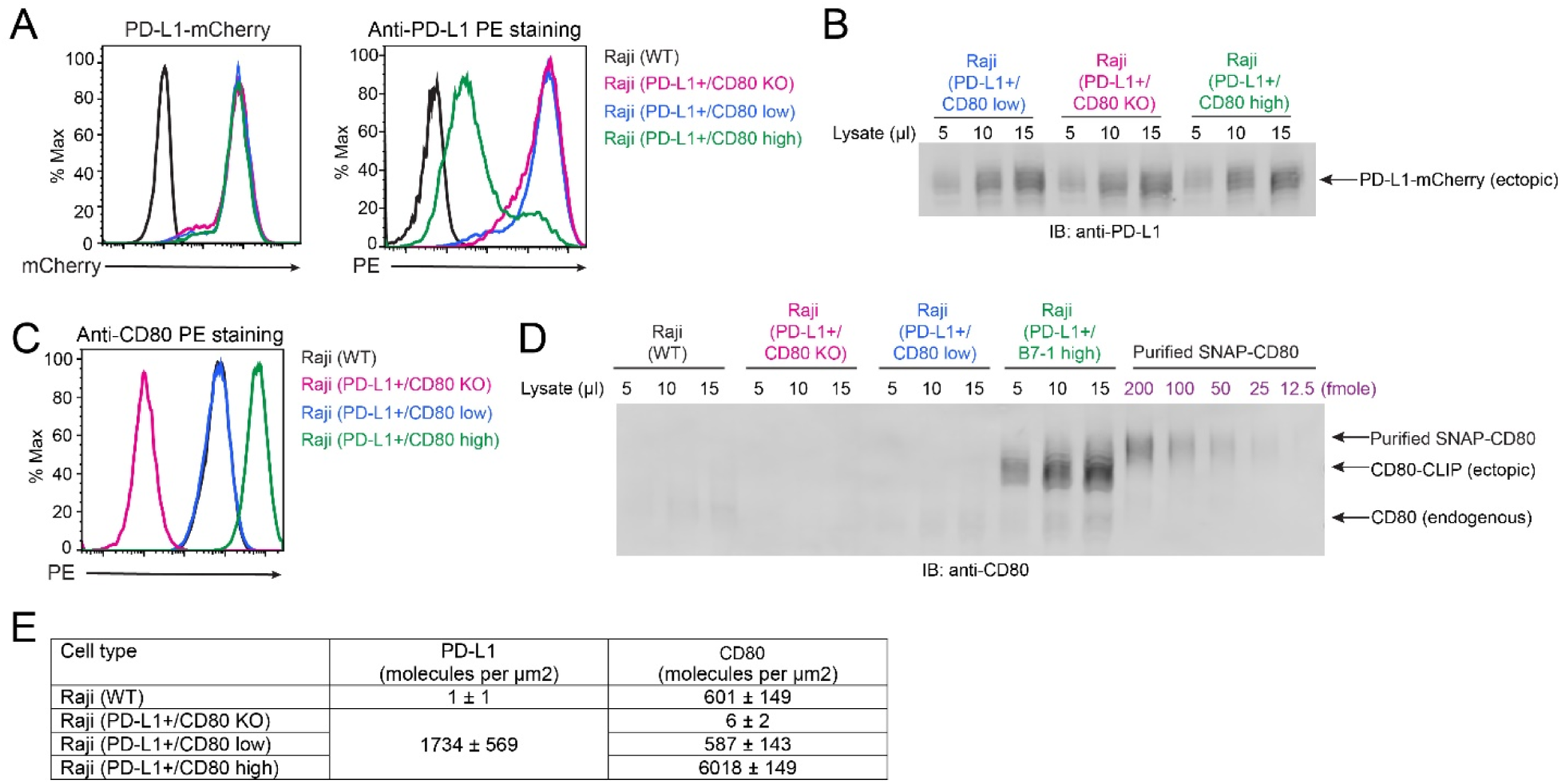
Quantification of PD-L1 and CD80 levels on Raji cell lines. Related to Figure 2. **(A)** (Left) Flow cytometry histograms of mCherry signals for the indicated unstained cell lines, reflecting the surface PD-L1-mCherry expression levels. (Right) Flow cytometry histograms of PE signals of the indicated cell lines stained with anti-PD-L1 PE (eBioscience, catalog 14-5983-82), reflecting the availabilities of anti-PD-L1 binding epitopes on the surface of each cell type. The much lower anti-PD-L1 staining for the Raji (PD-L1/CD80 high) cells suggests that PD-L1:CD80 cis-interaction blocks the anti-PD-L1 binding epitope. **(B)** Anti-PD-L1 (eBioscience, catalog 14-5983-82) immunoblots of the lysates of indicated cell lines, with the bands corresponding to the total PD-L1-mCherry expression levels. Cell lysates were treated with SDS sample buffer at 95 °C to disrupt PD-L1:CD80 *cis*-complexes. **(C)** Flow cytometry histograms of PE signals of the indicated cell lines stained with anti-CD80 PE (Biolegend, catalog 305208), reflecting the availabilities of anti-CD80 binding epitopes on the surface of each cell type. **(D)** Anti-CD80 (Novus Biologicals, NBP2-25255) immunoblots of the lysates of indicated cell lines, together with decreasing amounts of purified SNAP-CD80, from which a standard curve can be generated to calculate the CD80 levels in each cell line. **(E)** Table summarizing the PD-L1 and CD80 expression levels for the indicated cell lines. PD-L1 levels were calculated based on the mCherry signals in (**A**) and PD-L1-mCherry signals in (**B**), which agree with each other. The PE signals in (**A**) was not used because the PD-L1:CD80 cis-interaction blocks the anti-PD-L1 binding epitope. CD80 levels were calculated based on the PE signals in (**C**) and the anti-CD80 signals in (**D**), which agree with each other. Data are presented as mean ± SD, n = 3.

**Figure S2.**
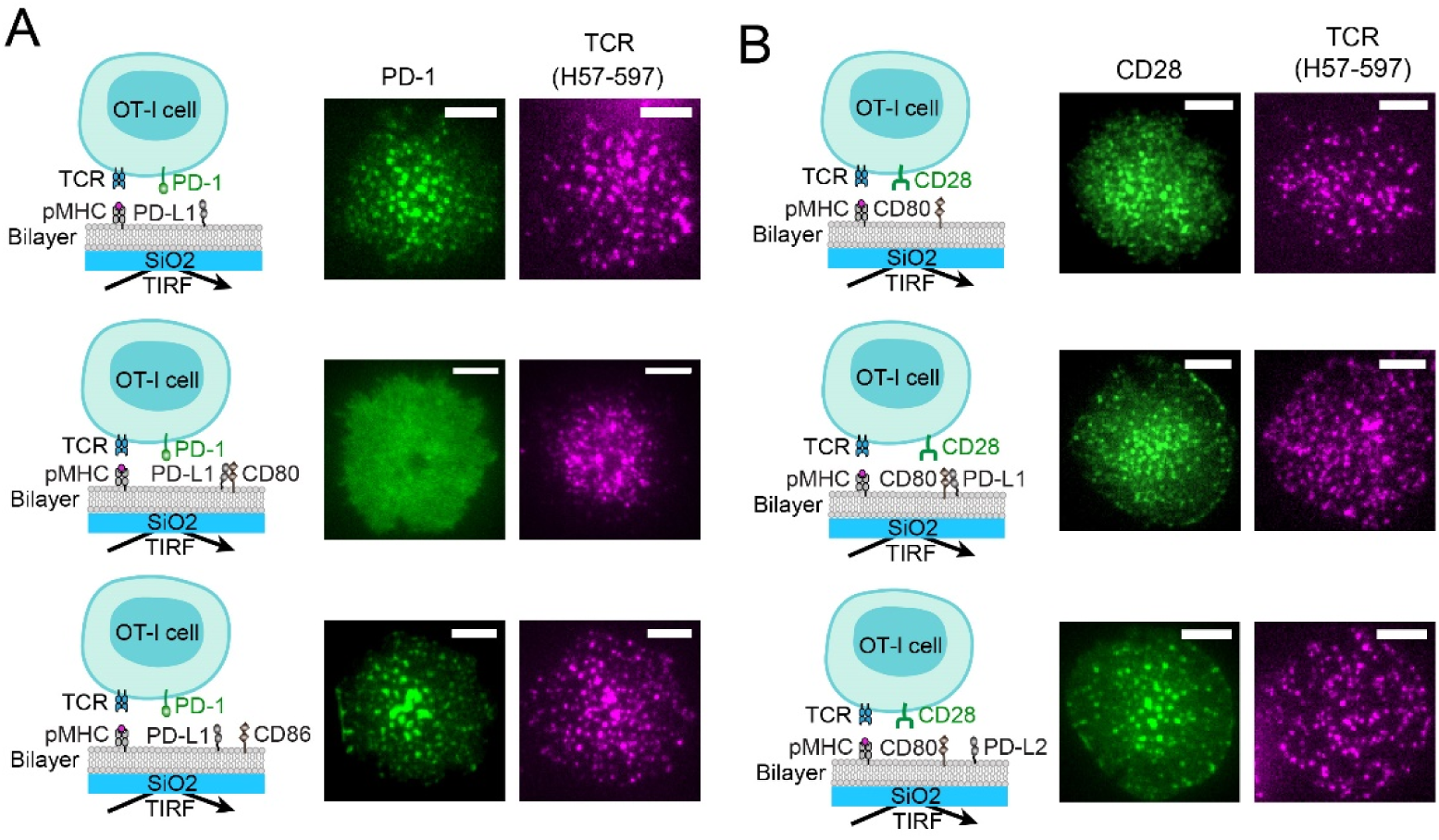
PD-L1:CD80 *cis*-interaction does not inhibit pMHC:TCR microclusters. Related to Figure 2 & Figure 3. **(A)** Cartoons on the left illustrate the experiment set up. PD-1-mCherry (rendered in green) transduced OT-I cells were stained with Alexa Fluor 647 labeled H57-597 TCR-β antibody, and plated onto SLB functionalized with pMHC/PD-L1 (top), pMHC/PD-L1 plus CD80 (three-fold excess to PD-L1), or pMHC/PD-L1 plus CD86 (three-fold excess to PD-L1). Shown next to each cartoon are representative TIRF images of synaptic PD-1 and TCR in a single OT-I cell 30 seconds after its initial contact with the SLB. **(B)** Cartoons on the left illustrate the experiment set up. CD28-mGFP (green) transduced OT-I cells were stained with Alexa Fluor 647 labeled H57-597 TCR-β antibody, and plated onto SLB functionalized with pMHC/CD80 (top), pMHC/CD80 plus PD-L1 (three-fold excess to CD80), or pMHC/CD80 plus PD-L2 (three-fold excess to CD80). Shown next to each cartoon are representative TIRF images of synaptic CD28 and TCR in a single OT-I cell 30 seconds after its contact with the SLB. Scale bars: 5 μm.

**Figure S3.**
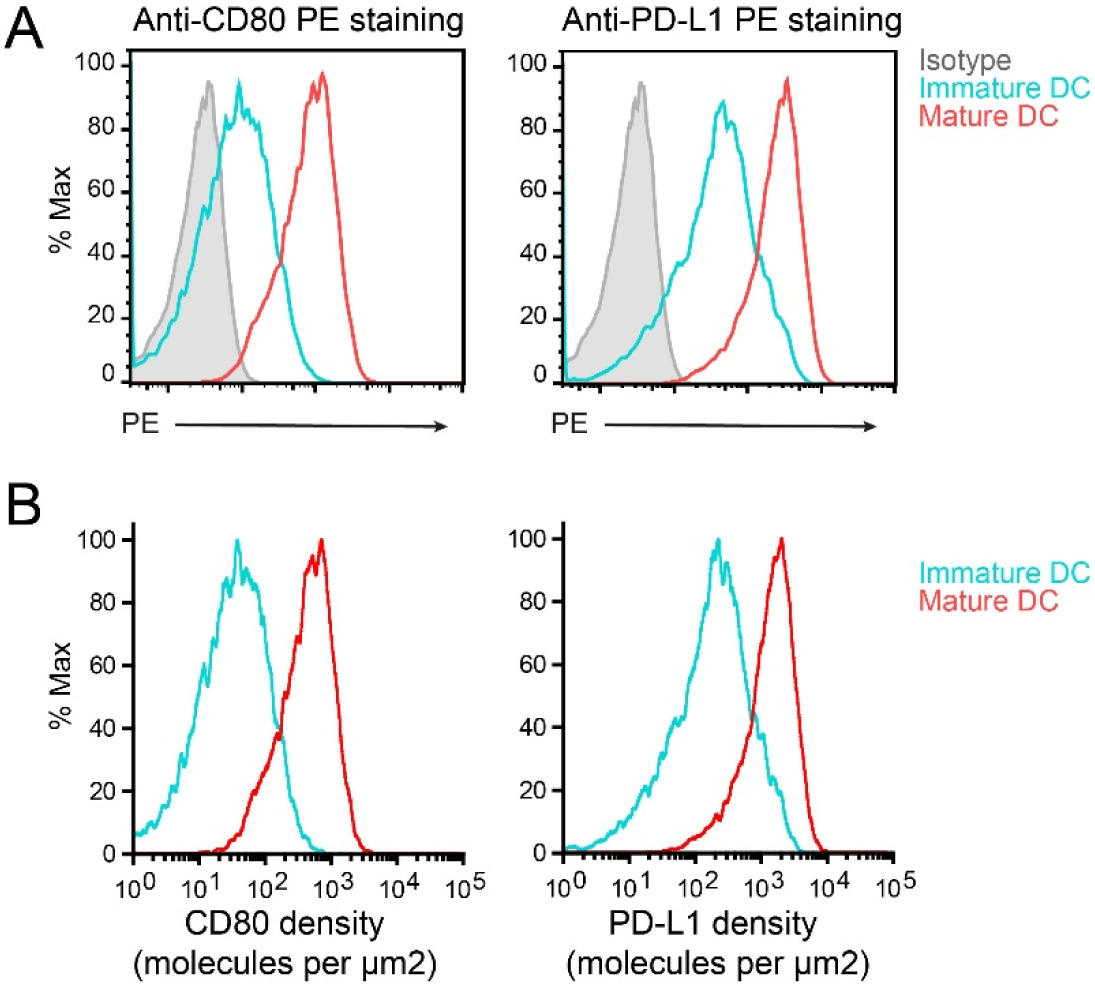
PD-L1 and CD80 expressions on human DCs. Related to Figure 2. **(A)** Flow cytometry histograms of PE signals on immature or mature human DCs stained with either anti-CD80 PE (Biolegend, catalog 305208) or anti-PD-L1 PE (eBioscience, catalog 14-5983-82). The grey histograms are the signal of isotype control under each condition. **(B)** Histograms of CD80 and PD-L1 surface densities on immature and mature DCs calculated from the flow cytometry histograms, using the QUANTUM™ R-PE MESF kit, as described in **Methods**.

**Figure S4.**
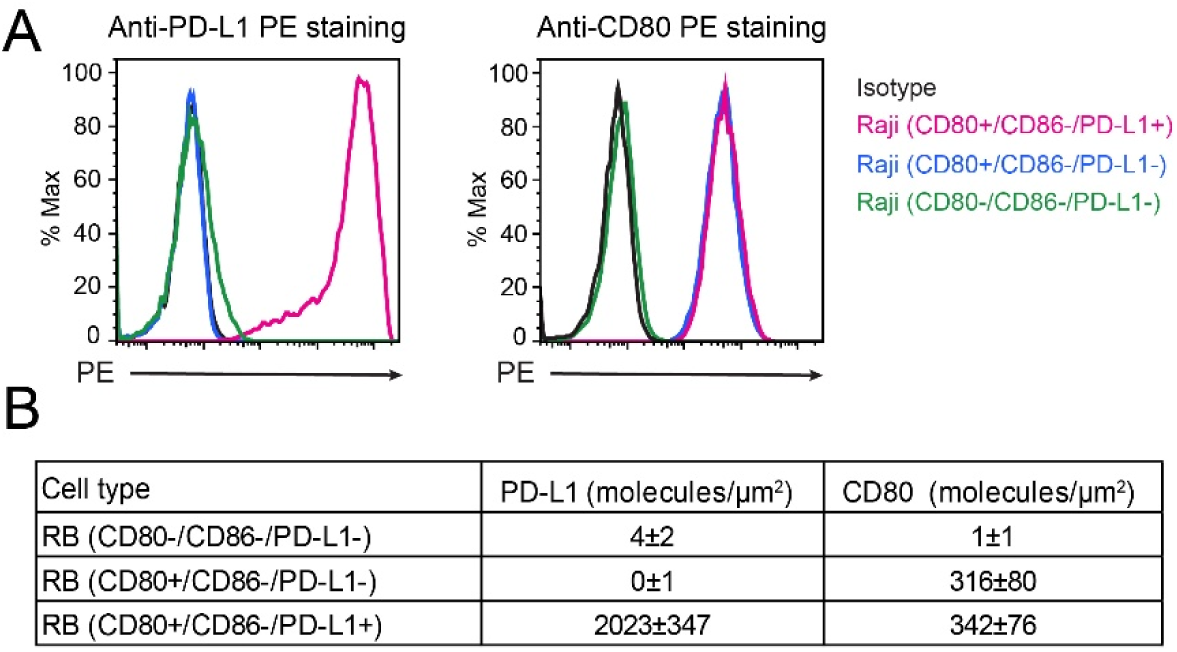
Quantification of PD-L1 and CD80 levels on Raji cell lines. Related to Figure 3 & Figure 4. **(A)** Flow cytometry histograms of PE signals of the indicated Raji cells stained with anti-PD-L1 PE (Left, eBioscience, catalog 14-5983-82) and anti-CD80 PE (Right, Biolegend, catalog 305208). **(B)** Table summarizing the PD-L1 and CD80 expression levels for the indicated cell lines. PD-L1 and CD80 levels were calculated based on PE signals in (**A**) using the QUANTUM™ R-PE MESF kit (Bangs Laboratories Inc, catalog 827). Data are presented as mean ± SD, n = 3.

